# MICAL2 acts through Arp3B isoform-specific Arp2/3 complexes to destabilize branched actin networks

**DOI:** 10.1101/2020.09.21.306522

**Authors:** Chiara Galloni, Davide Carra, Jasmine V. G. Abella, Svend Kjær, Pavithra Singaravelu, David J Barry, Naoko Kogata, Christophe Guérin, Laurent Blanchoin, Michael Way

## Abstract

The Arp2/3 complex (Arp2, Arp3 and ARPC1-5) is essential to generate branched actin filament networks for many cellular processes. Human Arp3, ARPC1 and ARPC5 exist as two isoforms but the functional properties of Arp2/3 iso-complexes is largely unexplored. Here we show that Arp3B, but not Arp3 is subject to regulation by the methionine monooxygenase MICAL2, which is recruited to branched actin networks by coronin-1C. Although Arp3 and Arp3B iso-complexes promote actin assembly equally efficiently in vitro, they have different cellular properties. Arp3B turns over significantly faster than Arp3 within the network and upon its depletion actin turnover decreases. Substitution of Arp3B Met293 by Thr, the corresponding residue in Arp3 increases actin network stability, and conversely, replacing Arp3 Thr293 with Gln to mimic Met oxidation promotes network disassembly. Thus, MICAL2 regulates a subset of Arp2/3 complexes to control branched actin network disassembly.

---

The Arp2/3 complex, consisting of seven subunits (Arp2, Arp3, ARPC1-5) is conserved in all eukaryotes, with the exception of some algae, protists and microsporidia (*1–4*). In mammals, the Arp3, ARPC1 and ARPC5 subunits exist as two isoforms (Arp3/3B, ARPC1A/B, and ARPC5/5L), which in humans are 91, 67 and 67% identical respectively (*5–7*). Several observations indicate that different Arp2/3 complexes have distinct regulatory or functional properties. For example, mutations that lead to a severe reduction or loss of ARPC1B expression in humans result in immunodeficiency and inflammation due to defects in cytotoxic T lymphocyte maintenance and activity (*8–13*). ARPC5L, together with g-actin, is required for peripheral nuclei positioning, while ArpC5 and β-actin are involved in transverse triad formation in *in vitro* assembled myofibers (*14*). The ARPC1 and ARPC5 isoforms also differentially affect the actin nucleating properties of the Arp2/3 complex as well as the stability of the branched filament networks they generate (*15*). In contrast, to ARPC1 and ARPC5 isoforms, the impact of Arp3B on the activity and function of the Arp2/3 complex remains to be established.

To investigate whether Arp3B impacts on the activity of the Arp2/3 complex we used actin-based motility of vaccinia virus as a model system (*15–18*). GFP-tagged Arp3 isoforms are both recruited to vaccinia-induced actin tails (Fig. 1A), consistent with their ability to incorporate into Arp2/3 complexes (Fig. S1A). Strikingly, GFP-tagged Arp3 and Arp3B induced the formation of longer and shorter actin tails respectively (Fig. 1A), suggesting that Arp3B may not be as effective as Arp3 in promoting actin polymerization. To investigate if this is the case, we examined the impact of RNAi-mediated depletion of Arp3 isoforms on actin-based motility of vaccinia (Fig. S1B). As described previously (*19, 20*), loss of Arp3 results in a significant reduction of all Arp2/3 complex subunits (Fig. 1B). Concomitant with this, there was a dramatic reduction in the length of vaccinia-induced actin tails (Fig. 1C, S1C), that is rescued by expression of GFP tagged Arp3 but not Arp3B (Fig. 1D, S1D, E). Knockdown of Arp3B had no appreciable impact on the level of Arp2/3 complex subunits (Fig. 1B). Nevertheless, there was a significant increase in actin tail length (Fig. 1C). Moreover, expression of GFP-Arp3B in cells treated with Arp3 siRNA results in short actin tails (Fig. 1D). These observations clearly demonstrate that Arp3 and Arp3B containing complexes have different ability to promote vaccinia-induced actin polymerization.

**Figure 1.**
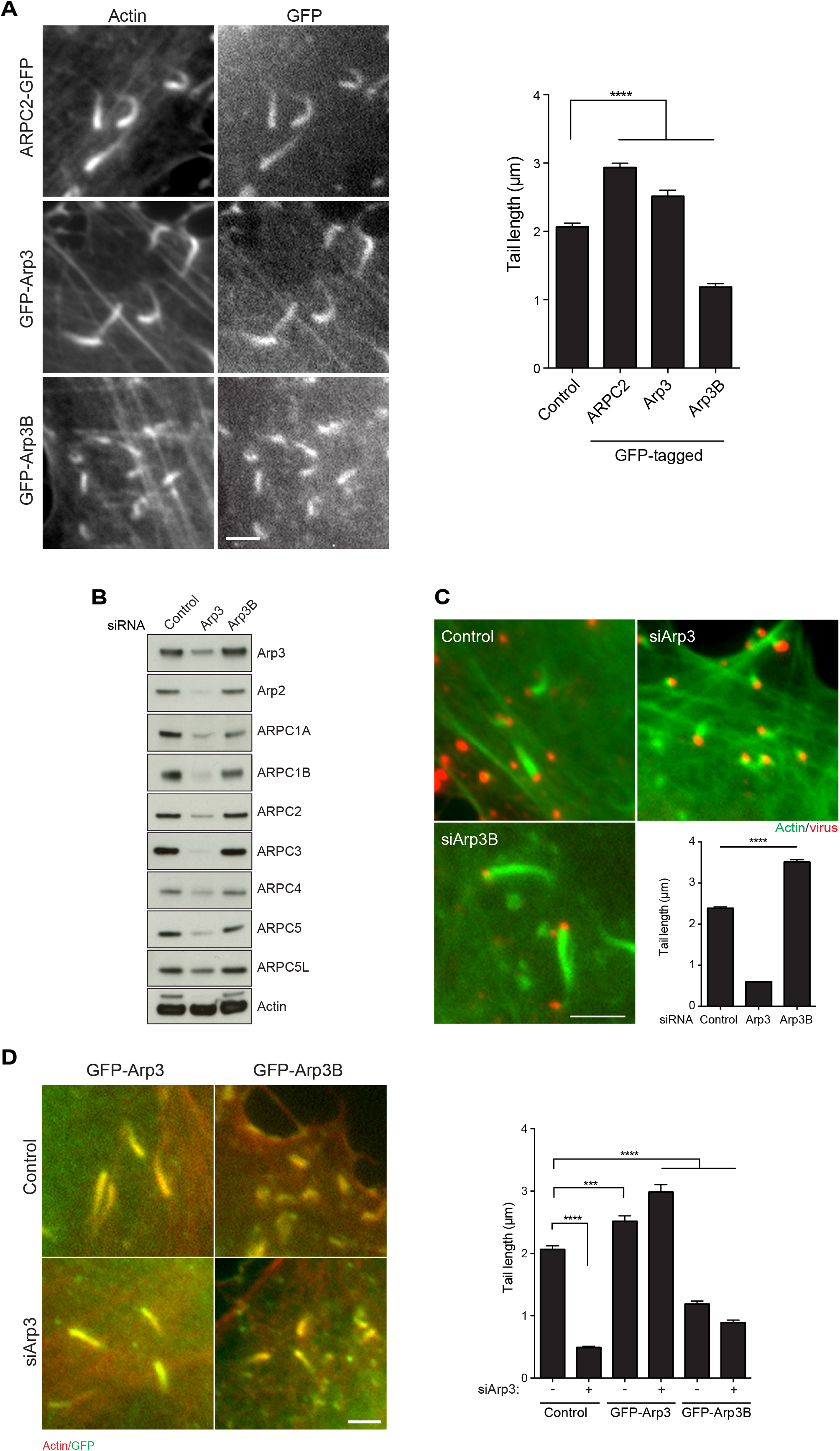
Arp3 isoforms confer different cellular properties to Arp2/3. **A** Localization of GFP tagged ARPC2, Arp3, or Arp3B to vaccinia-induced actin tails at 8 hours post-infection in HeLa cells. The graph shows the quantification of tail length relative to control HeLa cells. **B** Immunoblot analysis of lysates from HeLa cells treated with Arp3 and Arp3B siRNAs. **C** Immunofluorescence images of vaccinia (red) induced actin tails (green) together with quantification of their length in cells treated with the indicated siRNAs. **D** Images and quantification of actin tails (red) length in cells stably expressing RNAi resistant GFP-tagged Arp3 or Arp3B (green) and treated with Arp3 siRNA. All error bars represent SEM from three independent experiments in which 300 tails per condition were analyzed. **** P < 0.0001 and *** P < 0.001. Scale bar = 5μm.

To examine whether the differences in actin tail lengths are due to Arp3B complexes being less efficient than those with Arp3 in promoting actin polymerization, we performed in vitro pyrene actin assembly assays using defined recombinant human Arp2/3 complexes (Fig. 2A, S2A). We found that Arp2/3 complexes containing Arp3 and Arp3B were equally efficient at stimulating actin polymerization, with ARPC1B/ARPC5L containing complexes being better than those with ARPC1A/ARPC5 as observed previously (*15*) (Fig. 2A). We also performed TIRF microscopy to directly visualize Arp2/3 mediated actin assembly and branching (Fig. 2B, Movie S1). Automated quantification with the AnaMorf ImageJ plugin (*21*), reveals no differences in the rate of actin branch assembly (Fig. 2C). We therefore wondered whether short actin tails induced by Arp3B are due to increased actin disassembly. Using photoactivatable Cherry-GFP^PA^-actin we measured the rate of actin tail disassembly in cells treated with ARP3B siRNA (Fig. S2B). Loss of Arp3B increases the half-life of actin disassembly from 8.00 ± 0.16 to 9.82 ± 0.19 sec (Fig. 2D, S2C Movies S2 & S3), suggesting that Arp3B promotes the disassembly of branched actin networks. Furthermore, the halflife of Arp3B in actin tails (6.68 ± 0.17 sec) is significantly faster than that of Arp3 (8.63 ± 0.21 sec) (Fig. 2E, S2D, S2E Movies S4 & S5). Our data demonstrate that Arp3B containing Arp2/3 complexes generate less stable actin networks compared to those with Arp3.

**Figure 2.**
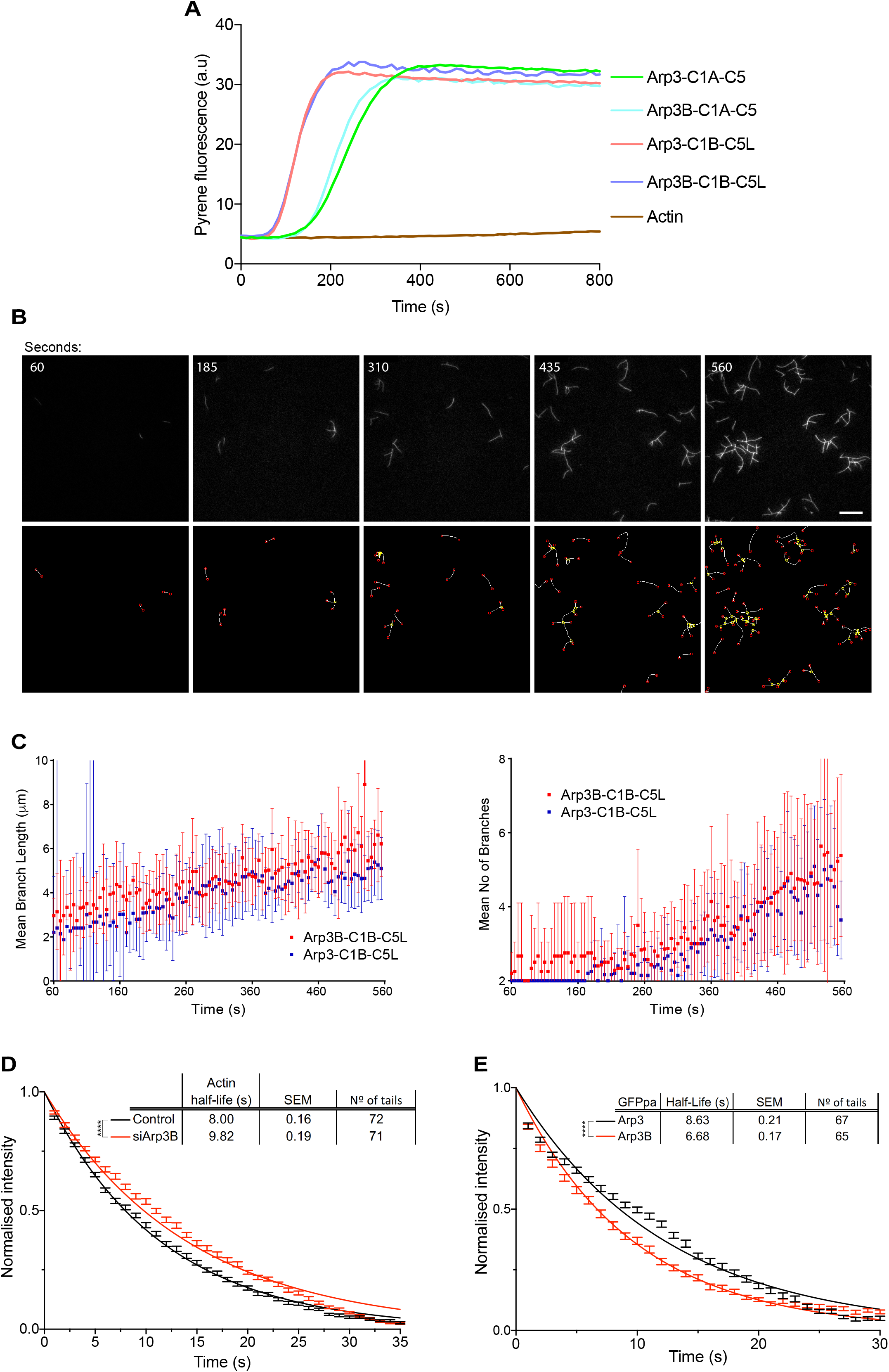
Arp3B reduces stability of actin networks. **A** In vitro pyrene actin assays using 12.5 nM of the indicated recombinant Arp2/3 reveal Arp3 and Arp3B containing complexes are equally efficient at nucleating actin regardless of the ARPC1A/ARPC5 and ARPC1B/ARPC5L background. **B** In vitro TIRF microscopy images to visualize branched actin formation. The top row shows image stills taken from TIRF assays using 2.5 nM Arp2/3 complexes containing Arp3B, ARPC1A and ARPC5 at the indicated times (see Movie S1) Scale bar = 15 μm. The bottom row shows the same panels after automatic detection of filament branches (yellow nodes) and ends (red nodes) with AnaMorf ImageJ plugin. **C** Quantification of mean branch number and length. **D** Quantification of the halflife of photoactivated GFP^PA^-β-actin in actin tails in Arp3B RNAi treated cells. These data were collected at same time as the data in Fig. 5G. **E** Quantification of the half-life of Arp3 and Arp3B in vaccinia actin tails after activation of the GFP^PA^ tag. The data in D and E from 3 independent experiments were combined and error bars represent SEM. for the indicated number of tails. Tukey’s multiple comparisons test was used to determine statistical significances, where **** P <0.0001.

Human Arp3 and Arp3B are 91% identical with amino acid differences spread throughout their length (Fig. S3A). To narrow down the region/amino acids responsible for the difference in actin network stability, we examined the impact of expressing a series of RNAi resistant Arp3/3B hybrids on actin tail length in cells lacking endogenous Arp3 and Arp3B (Fig. 3A, S3B). We found that residues 281-418 of Arp3B are sufficient to confer the short actin tail phenotype (Fig. 3B). This region contains 10 conservative and 3 non-conservative substitutions between the two proteins (Fig. S3A). Analysis of the structure of Arp2/3 in its active state reveals that two non-conserved substitutions (T293M and P295S) are on the surface of Arp3 in a hydrophobic loop which is positioned close to Arp2 (*22–24*) (Fig. 3C). To investigate their possible role in mediating isoform differences we exchanged residues 293 and 295 between Arp3 and Arp3B to generate GFP tagged Arp3^MS^ and Arp3B^TP^ (Fig. 3D). GFP-tagged Arp3^MS^ and ARP3B^TP^ switched phenotypes inducing the formation of short and long actin tails respectively in cells depleted of endogenous Arp3 and Arp3B (Fig. 3E, S3C, S3D). Actin tails remained long when single Arp3B substitutions were introduced into Arp3 (Fig. 3E, S3C, S3D). In contrast, actin tails remained short when Ser295 in Arp3B was changed to proline, but became long when Met293 was substituted for threonine. This demonstrates that Met293 is essential for Arp3B to induce short actin tails.

**Figure 3.**
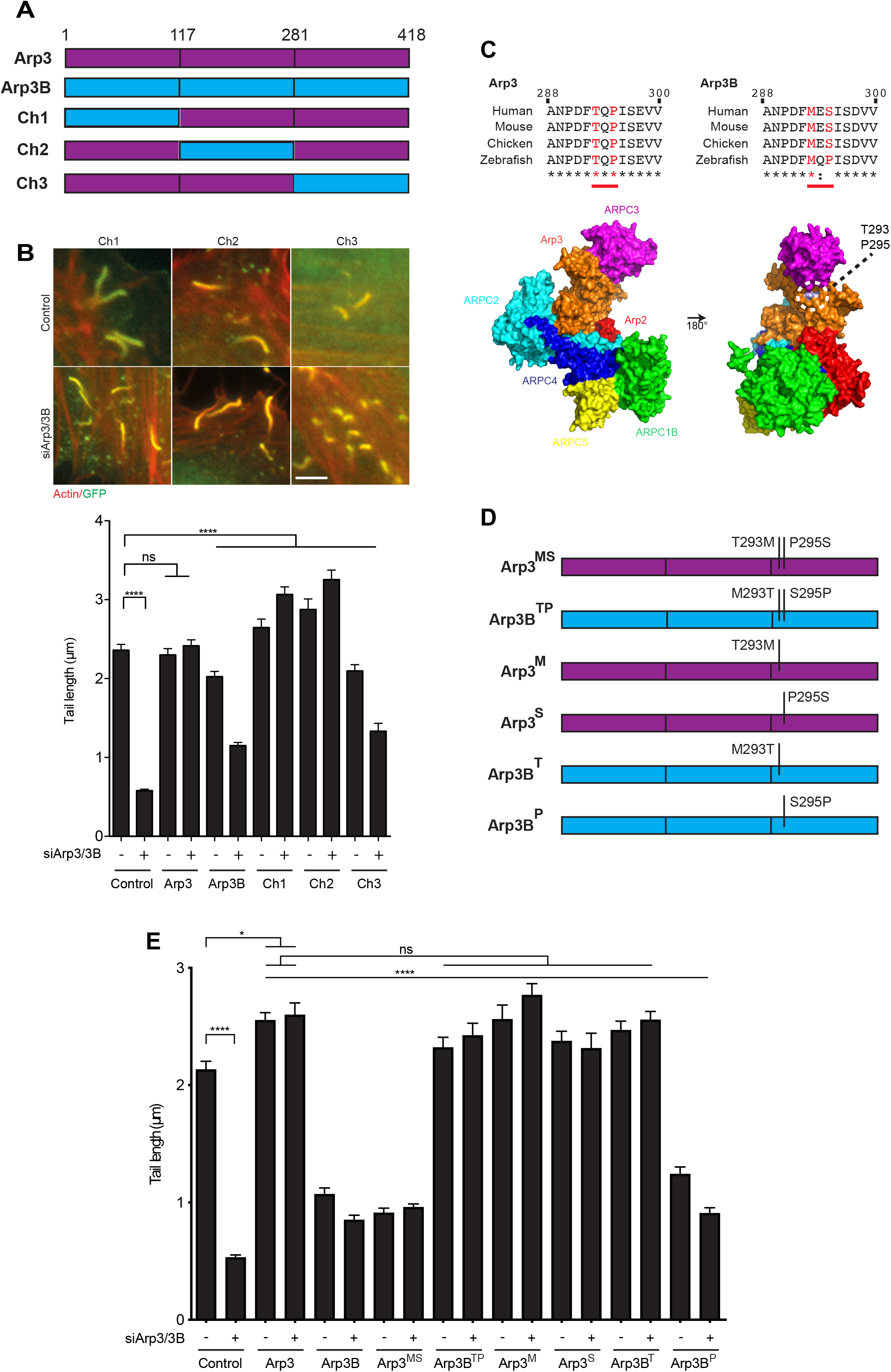
Met293 of Arp3B is essential to induce short actin tails. **A** Schematic representation of RNAi resistant human Arp3 and Arp3B together with the three different chimeras, Ch1, Ch2 and Ch3. Residue positions at the splice sites are indicated. **B** Images of actin tails (red) together with their quantification in HeLa cells stably expressing RNAi resistant GFP-tagged Ch1, Ch2 or Ch3 (green) and treated with Arp3 and Arp3B siRNA. Scale bar = 5 μm. **C** Alignment of residues 288-300 of Arp3 and Arp3B from the indicated species. Conserved residues are indicated with asterisks and colons highlight residues with similar properties. Residues 293 and 295 are shown in red. The structure highlights the position of Threonine 293 and Proline 295 on the surface of Arp3 in the bovine Arp2/3 complex (PDB 1K8K (*36*). **D** Schematic representation of the residue 293 and 295 exchanges between Arp3 and Arp3B. **E** Quantification of actin tail length in cells stably expressing RNAi resistant GFP-tagged protein and treated with Arp3 and Arp3B siRNA. All error bars represent SEM from three independent experiments in which 300 tails per condition were analyzed. **** P < 0.0001, * < 0.05 and ns is not significant.

Oxidation of Met44 and Met47 to methionine sulfoxide (Met-SO) in actin by MICAL proteins promotes actin filament disassembly (*25, 26*). Given this, we wondered whether the Arp3B-dependent actin tail phenotype depends on oxidation of Met293. To investigate this possibility, we replaced Thr293 in Arp3 and Met293 in Arp3B with glutamine to mimic the Met-SO state (*27, 28*). In both cases, the glutamine mutants induced the formation of shorter actin tails (Fig 4A, S4A). Consistent with the notion that Met293 in Arp3B is oxidized we found that GFP-tagged MICAL2 but not MICAL1 is recruited to actin tails (Fig. 4B, S4B Movies S6 & S7). Moreover, knockdown of MICAL2 also reduces the half-life of actin disassembly in tails by ~ 2 sec, while loss of MICAL1 had no impact (Fig. 4C, S4C). Depletion of MICAL2 or Arp3B alone or together resulted in similar long actin tails (Fig. 4D, S4D). Lastly, loss of MICAL2 suppressed the short actin phenotype induced by GFP-Arp3B over expression (Fig. 4E, S4E). Our data suggest that MICAL2 mediated oxidation of Met293 in Arp3B promotes faster actin network disassembly, resulting in shorter actin tails.

**Figure 4.**
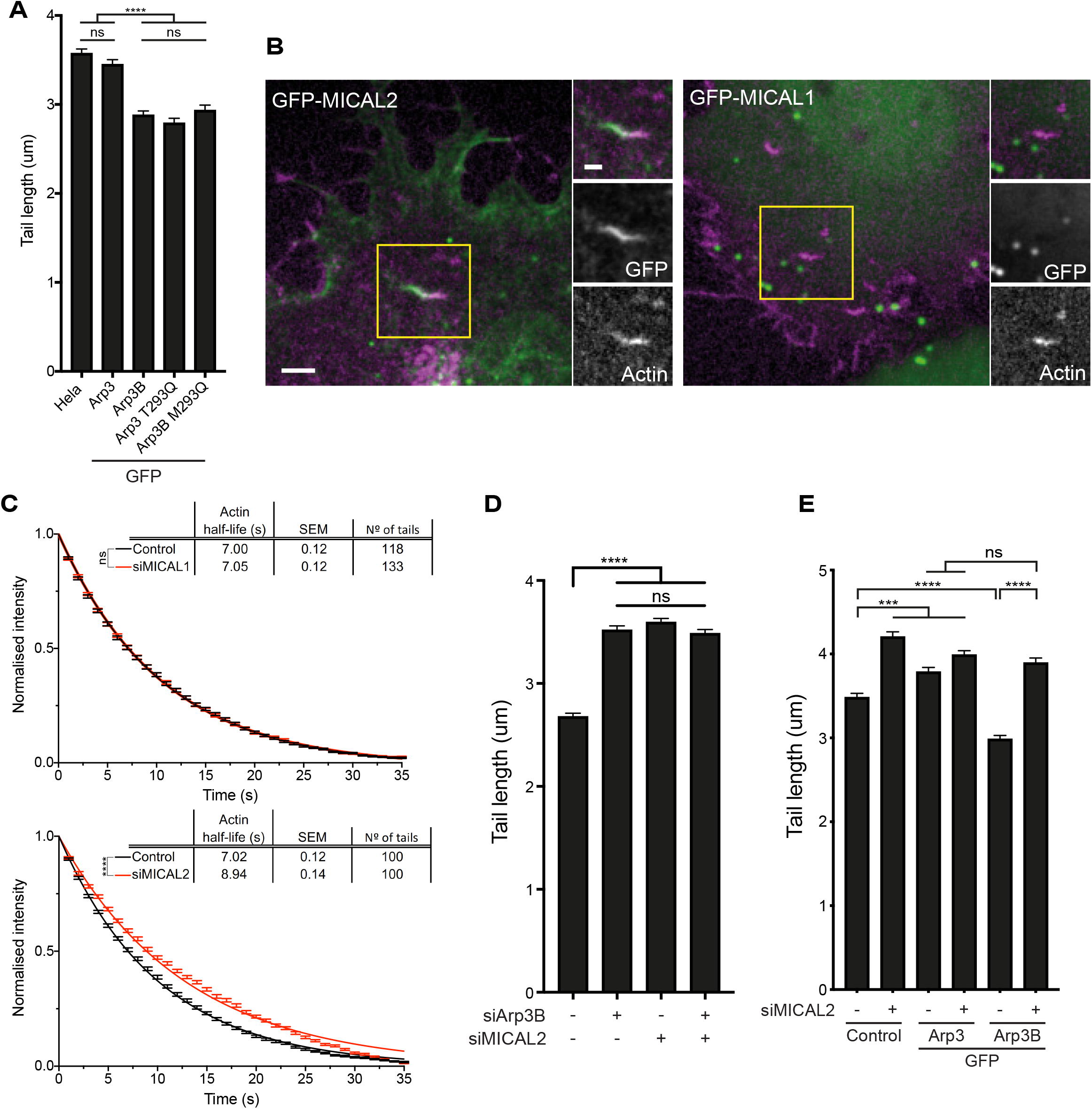
The ability of Arp3B to induce short tails depends on MICAL2. **A** Quantification of actin tail lengths in HeLa cells stably expressing the indicated GFP-tagged protein. **B** Image stills from Movie S6 and S7 showing that GFP-tagged (green) MICAL2 but not MICAL1 is recruited to Vaccinia induced actin tails (magenta). Scale bars = 5 μm (main image) and 2.5 μm (insert). **C** Quantification of the half-life of photoactivated Cherry-GFP^PA^-β-actin in actin tails in cells treated with MICAL1 and MICAL2 siRNA. The data from 3 independent experiments were combined and error bars represent SEM. for the indicated number of tails. Student’s t test was used to determine statistical significance. **D** The graph shows the length of actin tails in HeLa cells treated with the indicated siRNA. **E** Quantification of the effect of the loss of MICAL2 on actin tail length in HeLa cells stably expressing GFP-tagged Arp3 and Arp3B. All error bars represent SEM from three independent experiments in which 300 tails per condition were analyzed. Tukey’s multiple comparisons test was used to determine statistical significances, where **** P < 0.0001, *** < 0.001 and ns is not significant.

It is striking that GFP-MICAL2 is not recruited immediately behind the virus but further down the actin tail (Fig. 4B), as we previously observed for coronin (*15*). In light of this, we investigated if there is a connection between the two proteins given the role of coronin in promoting disassembly of branched actin networks (*29, 30*). Depletion of coronin-1C resulted in a dramatic loss of GFP-MICAL2 recruitment to actin tails (Fig. 5A, S5A, Movies S8 & S9) and suppression of the short actin tail phenotype induced by GFP-Arp3B (Fig. 5B, S5B). In agreement with this, GFP-Trap pulldowns demonstrated that coronin-1C associates with GFP-tagged MICAL2 but not MICAL1 (Fig. 5C). We previously demonstrated that coronin-induced actin tail disassembly depends on cortactin (*15*). Consistent with this, loss of cortactin also suppresses the short actin tail phenotype induced by GFP-Arp3B but had no impact on Arp3 induced actin tails (Fig. 5D, S5C). Furthermore, photoactivation experiments demonstrate that depletion of cortactin increases the half-life of Arp3B from 6.29 ± 0.17 to 9.48 ± 0.22 seconds but had no impact on Arp3 dynamics (Fig. 5E, S5D). In addition, depletion of coronin-1C or MICAL2 also results in a similar ~3 sec increase in Arp3B half-life, while having minimal impact on Arp3 (Fig. 5F, S5E). Concomitant with this, there was also a similar increase in the half-life of actin disassembly when Arp3B, coronin-1C or MICAL2 were depleted alone or in pairwise fashion (Fig. 5G, S5F).

**Figure 5.**
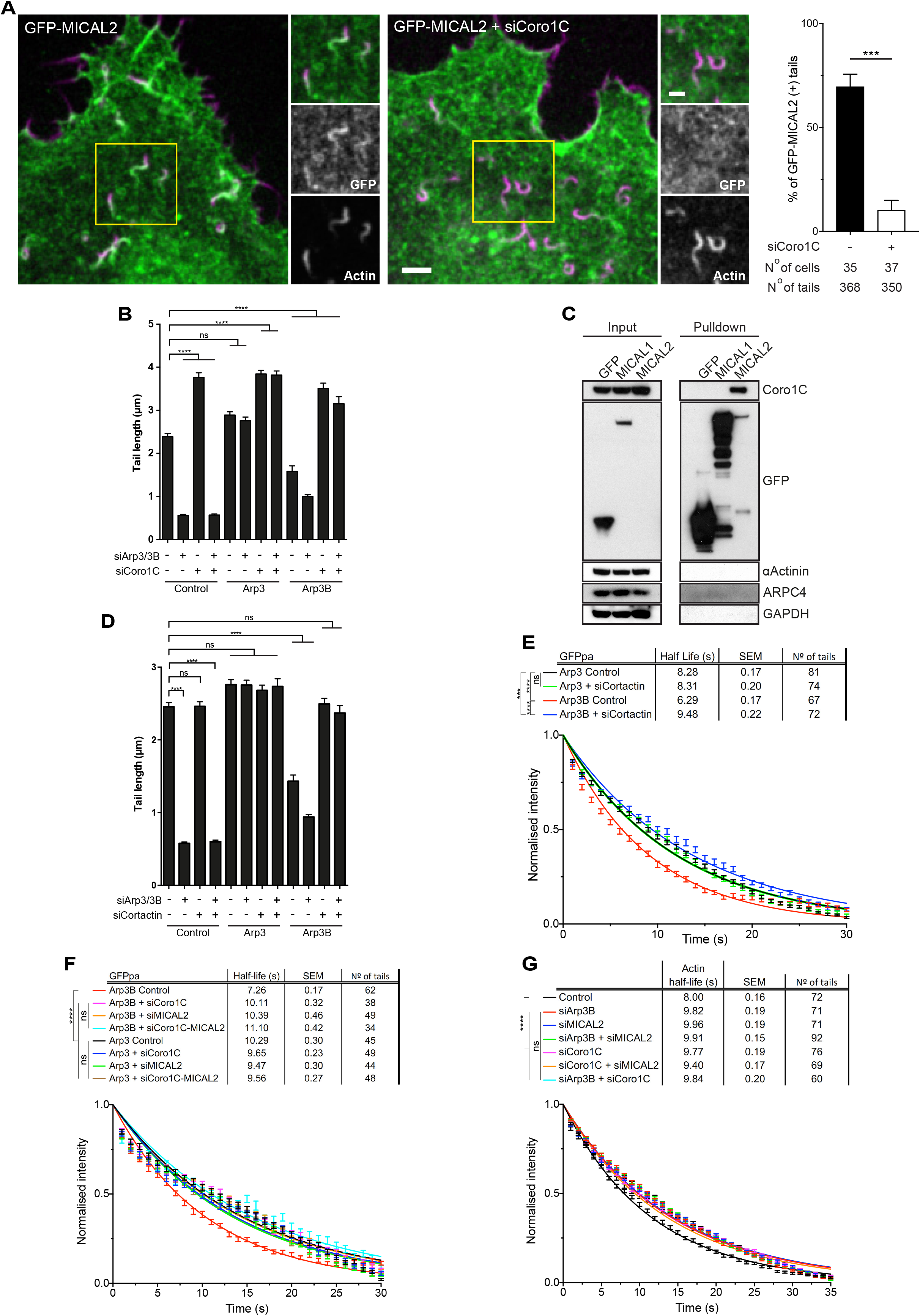
Coronin-1C recruits MICAL2 to actin networks. **A** Image stills from Movies S8 and S9 showing coronin-1C is required to recruit GFP-MICAL2 (green) to Vaccinia induced actin tails (magenta). Scale bars = 5 μm (main image) and 2.5 μm (insert). The graph shows the % of actin tails recruiting GFP-MICAL2 in HeLa cells following coronin-1C knockdown. The number of cells and actin tails analyzed is indicated. Student’s t test was used to determine statistical significance. **B** Quantification of actin tail length in cells stably expressing RNAi resistant GFP-tagged Arp3 or Arp3B and treated with Arp3/Arp3B and coronin-1C siRNA. All error bars represent SEM from three independent experiments in which 300 tails per condition were analyzed. **C** Immunoblot analysis of GFP-Trap pulldowns on HeLa cells stably expressing GFP-tagged MICAL1 or MICAL2. **D** Quantification of actin tail length in HeLa cells stably expressing RNAi resistant GFP-tagged Arp3 or Arp3B and treated with Arp3/Arp3B and cortactin siRNA. All error bars represent SEM from three independent experiments in which 300 tails per condition were analyzed. **E** Quantification of the half-life of photoactivated GFP^PA^-tagged Arp3 or Arp3B in actin tails in cells treated with cortactin siRNA. **F** Quantification of the half-life of photoactivated GFP^PA^-tagged Arp3 or Arp3B in actin tails in cells treated with coronin-1C and /or MICAL2 siRNA. **G** Quantification of the half-life of photoactivated Cherry-GFP^PA^-β-actin in actin tails in cells treated with different combinations of Arp3B, coronin-1C, MICAL2 siRNA. The data in E, F and G are from 3 independent experiments that were combined and error bars represent SEM. for the indicated number of tails. Tukey’s multiple comparisons test was used to determine statistical significances. **** P <0.0001, *** < 0.001 and ns is not significant

Actin assembly drives many cellular processes including cell migration, however, the disassembly of actin networks is equally as important for the actin cytoskeleton to perform its numerous functions (*15, 29, 30*). Our observations demonstrate that the recruitment of MICAL2 by coronin-1C enhances the disassembly of branched actin networks in an Arp3B but not Arp3 dependent fashion. Proteomic data indicates that Arp3B is widely expressed albeit at significantly lower levels than Arp3 (http://pax-db.org); for example, in HeLa cells Arp3B is 61 fold less abundant than Arp3 (*31*). Our data suggest that the relative levels of Arp3 and Arp3B will influence the stability of Arp2/3 induced branch networks. The regulation of a subset of Arp2/3 complexes by MICAL2, together with the impact of ARPC1 and ARPC5 isoforms, allows for further fine tuning of branched actin network dynamics in different cellular contexts and processes.

## Methods

### Vaccinia virus infection, antibodies, immunoblot and immunofluorescence analysis

HeLa were infected with the Western Reserve strain of Vaccinia virus 72 hours after siRNA transfection and processed for immunofluorescence analysis as previously described (*15*). Vaccinia was detected with a monoclonal antibody against B5 (19C2) (1:500 dilution) (*32*). Actin tails were labelled with Alexa488 or Texas red phalloidin (Invitrogen). Samples from each siRNA condition were kept for immunoblot analysis. All secondary (Alexa conjugated) antibodies were purchased from Molecular Probes, Invitrogen.

The other antibodies used in this study are listed below: ARPC2/p34-Arc polyclonal (Millipore, 07-227, 1:1000 dilution), ARPC5 monoclonal (Synaptic Systems, 305011, clone 323H3, 1:1000 dilution), Arp2 monoclonal (Abcam, ab129018, clone EPR7979, 1:1000 dilution), ARPC5L monoclonal (Abcam, ab169763, clone EPR10274, 1:1000 dilution), ARPC1A polyclonal (Sigma, HPA004334, 1:250 dilution), Arp3 monoclonal (Sigma, A5979, clone FMS338, 1:1000 dilution), ARPC1B polyclonal (Bethyl Laboratories, A302-781, 1:1000 dilution), ARPC3 monoclonal (BD Transduction labs, 612234, clone 26/p21-Arc, 1:1000 dilution), ARPC4 polyclonal (Sigma, SAB1100901, 1:1000 dilution), β-actin monoclonal (Sigma, A5316, clone AC-74, 1:1000 dilution), Vinculin monoclonal (Sigma, V4505, 1:1000 dilution), coronin-1C polyclonal (Invitrogen, PA5-21775, 1:500 dilution), a-Actinin monoclonal (Sigma, A5044, 1:1000 dilution), GAPDH monoclonal (Santa Cruz, sc-32233, 1:1000 dilution), Cortactin monoclonal (Millipore, 05-180, clone 4F11, 1:1000 dilution), MICAL1 polyclonal (Proteintech, 14818-1-AP, 1:1000 dilution), GFP monoclonal (custom made by CRUK, 1:1000 dilution).

### Stable cell lines

The Cherry–GFP^PA^-β-actin stable HeLa cells were previously described (*15*). Stable HeLa cell lines expressing siRES GFP-Arp3, GFP-Arp3B, GFP-Ch1, GFP-Ch2, GFP-Ch3, GFP-Arp3^MS^, GFP-Arp3B^TP^, GFP-Arp3^M^, GFP-Arp3^S^, GFP-Arp3B^T^, GFP-Arp3B^P^, GFP-MICAL1 and GFP-MICAL2 were generated using the lentivirus infection (Trono group second generation packaging system, Addgene) and selected using puromycin resistance (1 μg/ml)) as previous described (*15*). The siRES GFP^PA^-Arp3 or GFP^PA^-Arp3B lentiviral constructs were used to infect Lifeact-RFP stable HeLa cells (*15*) and cells expressing both proteins were selected using a combination of hygromycin and puromicin (100 μg/ml and 1 μg/ml respectively).

### Expression constructs

The expression vectors pLVX-LifeAct-RFP, pLVX-GFP^pa^-Cherry-β-actin, pLVX-ARPC2-GFP have been previously described (*15*). Human Arp3, Arp3B, Chimera 1, 2, 3, Arp3^MS^, Arp3B^TP^ and the single point mutants were obtained as synthetic genes (Geneart, Invitrogen) and cloned into the NotI/EcoRI sites of pLVX-N-term-GFP (*15*). Using pLVX-GFP-Arp3 and pLVX-GFP-Arp3B as a template, primers containing a mismatch for the T293Q and M293Q mutation were used to amplify two overlapping fragments corresponding to Arp3 or Arp3B respectively. These two fragments were then assembled into NotI/EcoRI digested pLVX-N-term-GFP using Gibson-Assembly to obtain pLVX-GFP-Arp3-T293Q and pLVX-GFP-Arp3B-M293Q. Human MICAL1 was obtained by PCR amplification from a GFP-MICAL1 plasmid provided by Arnaud Echard (Institut Pasteur, Paris) (*33*). This PCR fragment was cloned into the NotI/EcoRI sites of pLVX-N-term-GFP to generate pLVX-GFP-human-MICAL1. Human MICAL2 was obtained by PCR amplification from a HA-MICAL2 plasmid donated by Steve Caplan (University of Nebraska, USA) (*34*). This PCR fragment was cloned into the NotI/EcoRI sites of pLVX-N-term-GFP using Gibson-Assembly to generate pLVX-GFP-human-MICAL2. LifeAct-iRFP670 was amplified from pLVX-LifeAct-iRFP670 (Snetkov et al. 2016) and cloned using Gibson Assembly into a BglII/BamHI digested pE/L vector to obtain pE/L-LifeAct-iRFP670.

### Live cell imaging and photoactivation

Photoactivation experiments involving GFP^PA^-tags were performed as previously described (*15*). To visualise actin structures in live imaging, GFP-MICAL1 and GFP-MICAL2 stable cells were transfected with pE/L-LifeAct-iRFP670 using FUGENE (Promega) and infected with vaccinia virus as previously described (*35*). Three-minute videos were acquired 8 hours post infection on an Axio Observer Microscope controlled by Slidebook (3i intelligent imaging innovations). Videos of Control and siCoro1C transfected cells were pooled and blind scored for the presence of GFP-MICAL2 on actin tails using ImageJ. Data are presented as the mean ± standard error of the mean of the percentage of GFP-positive tails over four independent experiments.

### siRNA transfection

HeLa cells were transfected with siRNA as previously described (*15*). The following siRNA were used: All-Star control (Qiagen, SI03650318), siRNA against Arp3 (Dharmacon M-012077-01-0010), Arp3B (Dharmacon M-020372-01-0010), Cortactin (Dharmacon MU-010508-00-0010), Coronin-1C (Dharmacon M-017331-00-0010) and MICAL2 (Dharmacon L-010189-00-0005). A MICAL1 targeting siRNA (5’-GAGUCCACGUCUCCGAUUU-3) was designed as previously described (*33*). The following deconvoluted siRNA were used: against Arp3 (Dharmacon D-O12077-01, D-O12077-03, D-O12077-04, D-O12077-05) and Arp3B (Dharmacon D-020372-01, D-020372-02, D-020372-03, D-020372-04).

### GFP-Trap pulldown assays

Cells were treated with 200 nm LatA for 1h and then lysed in the following buffer: 50 mM HEPES, pH 7.5, 150 mM NaCl, 1.5mM MgCl_2_, 1mM EGTA, 1% Triton X-100, 10% glycerol, 50mM NaF, 1mM sodium vanadate, 200 nM LatA, Phosphostop and protease inhibitors (Roche). Lysates were normalised and incubated with GFP-Trap agarose beads (Chromotek) and pulldown was performed according to manufacturer’s instructions.

### Recombinant expression and purification of Arp2/3 complexes

Human Arp3B was amplified by PCR using pLVX-GFP-Arp3B as a template and cloned into the BamHI/NotI sites of the pFL vector which also contains ArpC2 (*15*). The other baculovirus expression vectors and the Arp2/3 complex purification protocol were previously described (*15*).

### Pyrene actin polymerisation, TIRF branching assays and analysis

Pyrene labelled and unlabelled actin monomers were mixed in G-buffer (2 mM Tris (pH 8.0), 0.2 mM ATP, 0.1 mM CaCl2, and 0.5 mM DTT) to obtain the desired ratio of pyrene-actin to unlabelled actin (10 % pyrene-actin), and diluted to 20 μM total actin in G-buffer. Actin polymerization was performed in presence of 2 μM of monomeric actin (10% pyrene labeled) at room temperature by the addition of one-tenth volume of 10X KMEI (5 mM Tris-HCl [pH 8.0], 0.2 mM ATP, 0.1 mM CaCl2, and 0.5 mM DTT) in G-buffer. Experiments were conducted in the presence or in the absence of different Arp2/3 complexes. The polymerization was followed by fluorescence measurements with excitation and emission wavelengths of 365 nm and 407 nm respectively using a Xenius SAFAS fluorimeter (Safas SA, Monaco). The data were analyzed with Graphpad Prism 6.

Time course of actin assembly (0.6 μM) in presence of 2.5 nM Arp2/3 complex and 50 nM GST-(Human WASp-VCA) was acquired on a total internal reflection fluorescence (TIRF) microscope (Roper Scientific) equipped with an iLasPulsed system and an Evolve camera (EMCCD) using a 60X 1.49 numerical aperture (NA) objective lens. Microscope and devices were driven by MetaMorph (Molecular Devices). Automated quantification of TIRF actin assembly assays was performed with the AnaMorf ImageJ plugin (*21*) details available at https://github.com/djpbarry/AnaMorf/wiki.

### Reverse Transcription Quantitative PCR (RT-qPCR)

RNA extraction, cDNA synthesis and qPCR were performed as previously described (*35*). The following primers were used: GAPDH forward TGATGACATCAAGAAGGTGGTG, reverse TCCTTGGAGGCCATGTGGGCCA; ACTR3 forward CGATATGCAGTTTGGTTTGG, TTTGGTGTGGCATACTTGGT; ACTR3B forward GCCCGCTGTATAAGAATGTCG, reverse AATCCCTGAACATGGTGGAGC; MICAL2 forward GTGCACGAACACCAAGTGTC, reverse CCAGAGGTGTAGCACGTTGT; Coronin-1C forward GGTTTCTCGTGTGACCTGGG; reverse TCAATTCGACCAGTCTTGTGC.

### Statistical Analysis

For actin tail length experiments, 10 cells per condition were acquired and 10 tails per cell were measured in each independent experiment (100 tails per condition in total). Data from three independent experiments were combined and the error bars represent the SEM for n = 300 tails per condition. When only 2 conditions were compared, Student t test was performed to determine statistical significance. When more than two samples were compared, Tukey’s multiple comparisons test was used to determine statistical significances. All data concerning was analysed using Graphpad Prism 8.

## Acknowledgements

We would like to thank Arnaud Echard (Institut Pasteur, Paris) and Steve Caplan (University of Nebraska, USA) for providing GFP-MICAL1 and HA-MICAL2 plasmids respectively. Joseph Cockburn (Leeds University) and Carolyn Moores (Birkbeck College, London) for structural insights. Alessio Yang and Angika Basant from Way lab for providing the pE/L-LifeAct-iRFP670 expression vector. We would like to thank Richard Treisman and Angika Basant (Francis Crick Institute) for critically reading the manuscript and scientific discussion.

## Author contributions

CG, DC and NK generated stable HeLa cell lines (GFP-tagged Arp3, Arp3B, Ch1, Ch2, Ch3, Arp3^MS^, Arp3B^TP^, Arp3^M^, Arp3^S^, Arp3B^T^, Arp3B^P^), (GFP-tagged Arp3-T293Q, Arp3B-M293Q, MICAL1, MICAL2) and (GFP^PA^-Arp3 and GFP^PA^-Arp3B) respectively. Actin tail lengths were quantified by CG (Figs. 1, S1, 3, 5) and DC (Fig. 4). Immunoblots and RT-PCR were generated by CG (Figs. 1, S1, S2A, S3, S5B and S5C) and DC (Figs. S2B, D, S4, 5C, S5A, S5D, S5E and S5F). GFP-TRAP pulldowns in Figs. S1A and 5C were performed by CG and DC respectively. All photoactivation experiments (Figs. 2D, 2E, 4C, 5E, 5F, and 5G) and their analyses were performed by DC, as were GFP-MICAL localization experiments (Figs. 4B and 5A). SK expressed and purified recombinant Arp2/3 complexes. PS, C. Guérin and LB performed actin assembly assays and provided scientific discussion. DJB analysed TIRF assays and provided advice concerning analysis of photoactivation experiments. JGVA and MW conceived and supervised the work.

## Supplementary figure legends

**Figure S1.**
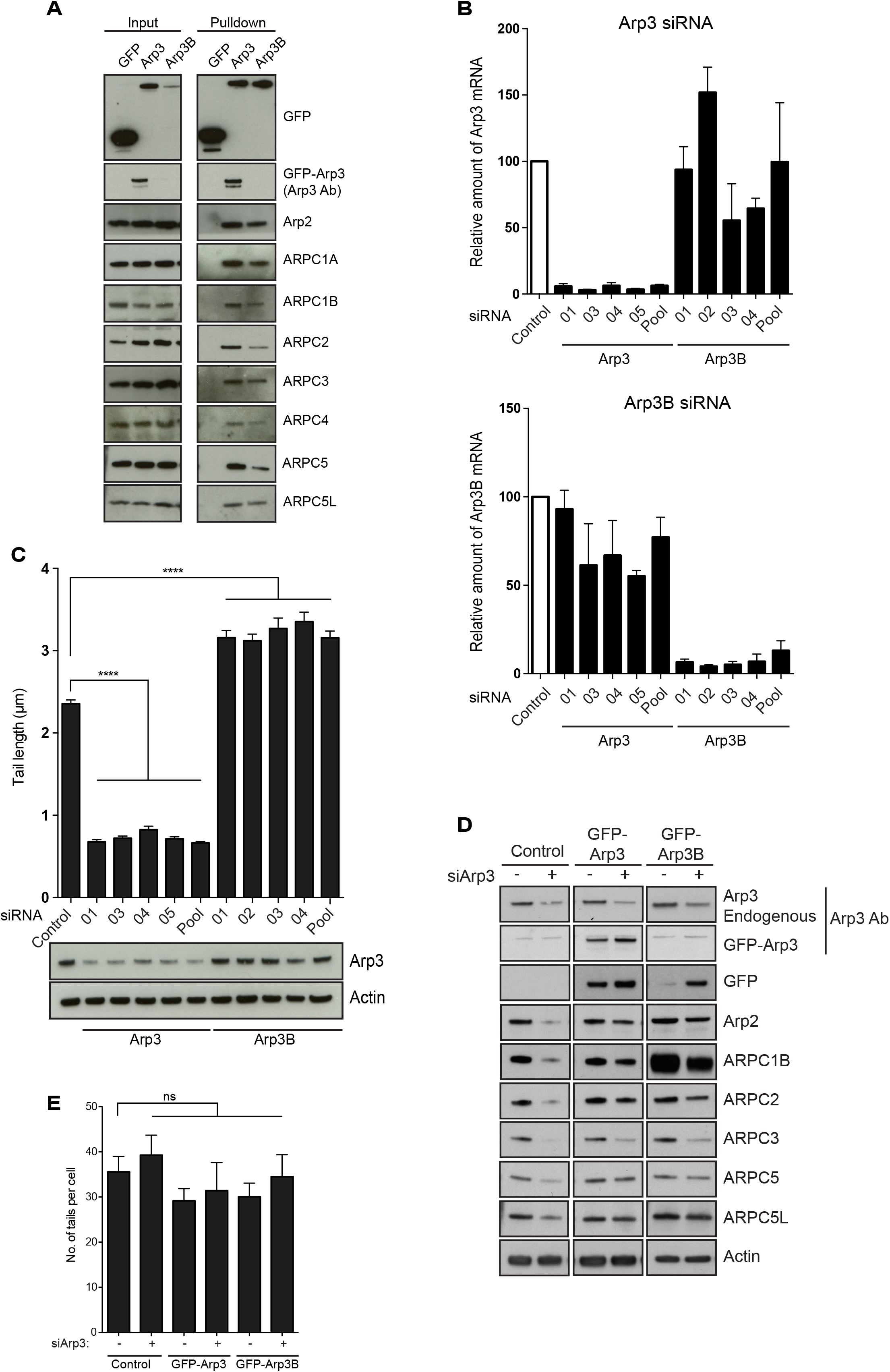
Arp3 and Arp3B knockdown and rescue experiments. **A** Immunoblot analysis of GFP-Trap pulldowns from HeLa cells lysates stably expressing GFP, GFP-Arp3 or GFP-Arp3B demonstrates that the 8 different isoform combinations are formed. **B** RT-PCR analysis showing the level of Arp3 and Arp3B mRNA relative to control in HeLa cells after knockdown with the individual siRNA from the siGenome pools against Arp3 and Arp3B used in Fig. 1. **C** Quantification of actin tail lengths and immunoblot analysis of the level of Arp3 in HeLa cells treated with the indicated siRNA. **D** Immunoblot analysis showing that stable expression of RNAi resistant GFP-tagged Arp3 and Arp3B restore control levels of all Arp2/3 subunits in HeLa cells treated with Arp3 siRNA. **E** Quantification of the number of actin tails in cells stably expressing RNAi resistant GFP-tagged Arp3 or Arp3B and treated with Arp3 siRNA. Error bars represent SEM from three independent experiments in which 300 tails per condition were analyzed. **** P < 0.0001 and ns is not significant.

**Figure S2.**
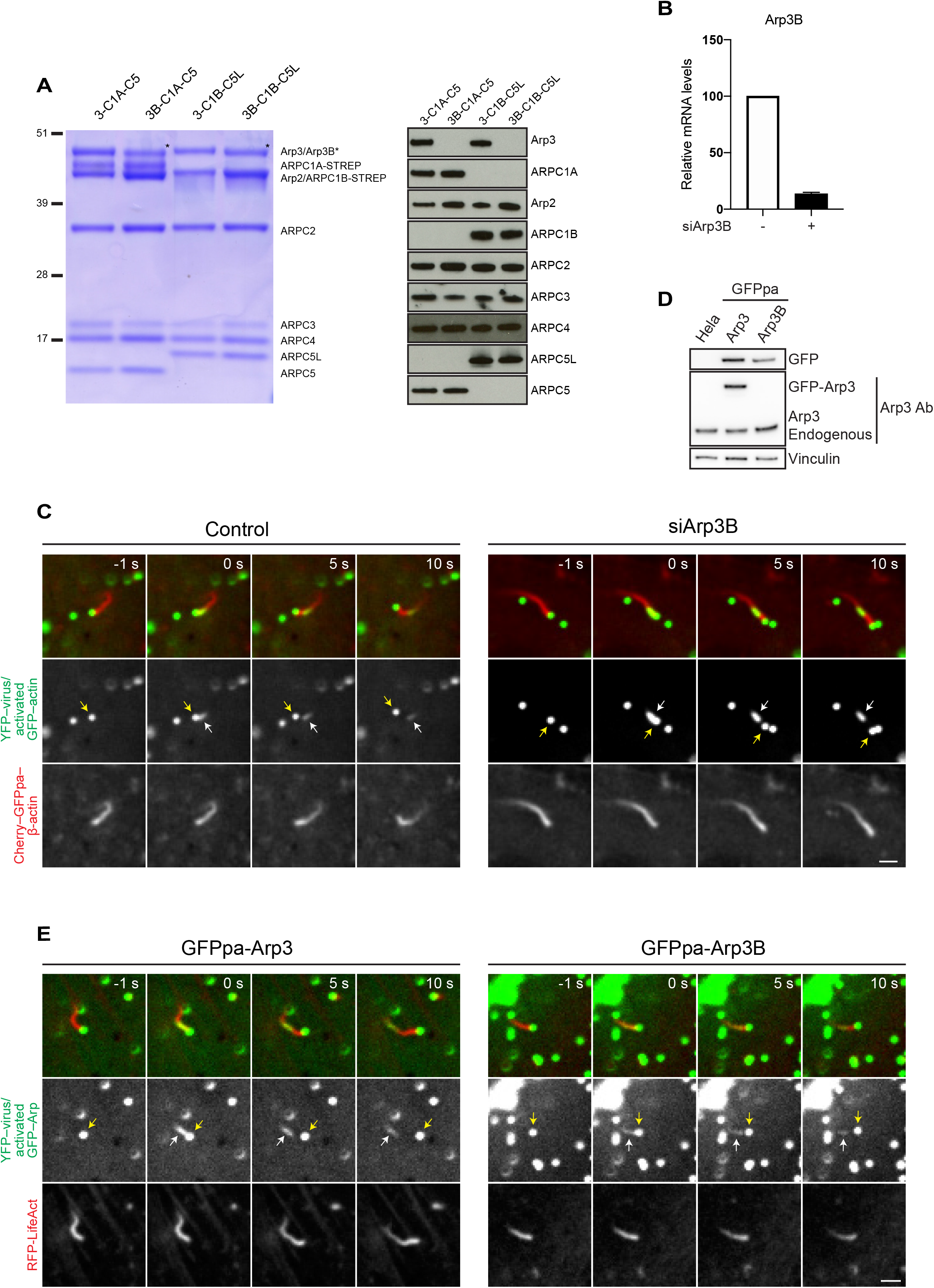
Arp3 and Arp3B dynamics. **A** Coomassie-stained gel of recombinant isoform-specific Arp2/3 complexes and corresponding immunoblot. **B** RT-PCR analysis of the level of Arp3B mRNA in HeLa cells stably expressing Cherry-GFP^PA^-β-actin and treated with Arp3B siRNA. Error bars represent SEM from three independent experiments. **C** Image stills from Movie S2 and Movie S3 at the indicated times in seconds before and after photoactivation showing the movement of YFP-tagged virus (yellow arrow) in HeLa cells stably expressing Cherry-GFP^PA^-β-actin (red before photoactivation) in control and Arp3B RNAi treated cells. The white arrow indicates the position of photoactivated Cherry-GFP^PA^-β-actin at time zero (green). **D** Immunoblot analysis of lysates from HeLa cells stably expressing GFP^PA^-tagged Arp3 and Arp3B. **E** Image stills from Movie S4 and Movie S5 at the indicated times in seconds showing the movement of YFP-tagged virus (yellow arrow) in HeLa cells stably expressing LifeAct-RFP together with GFP^PA^-tagged Arp3 or Arp3B. White arrow indicates the position of photoactivation of GFP^PA^-tagged Arp3 or Arp3B at time zero (green). Scale bars 2.5 μm.

**Figure S3.**
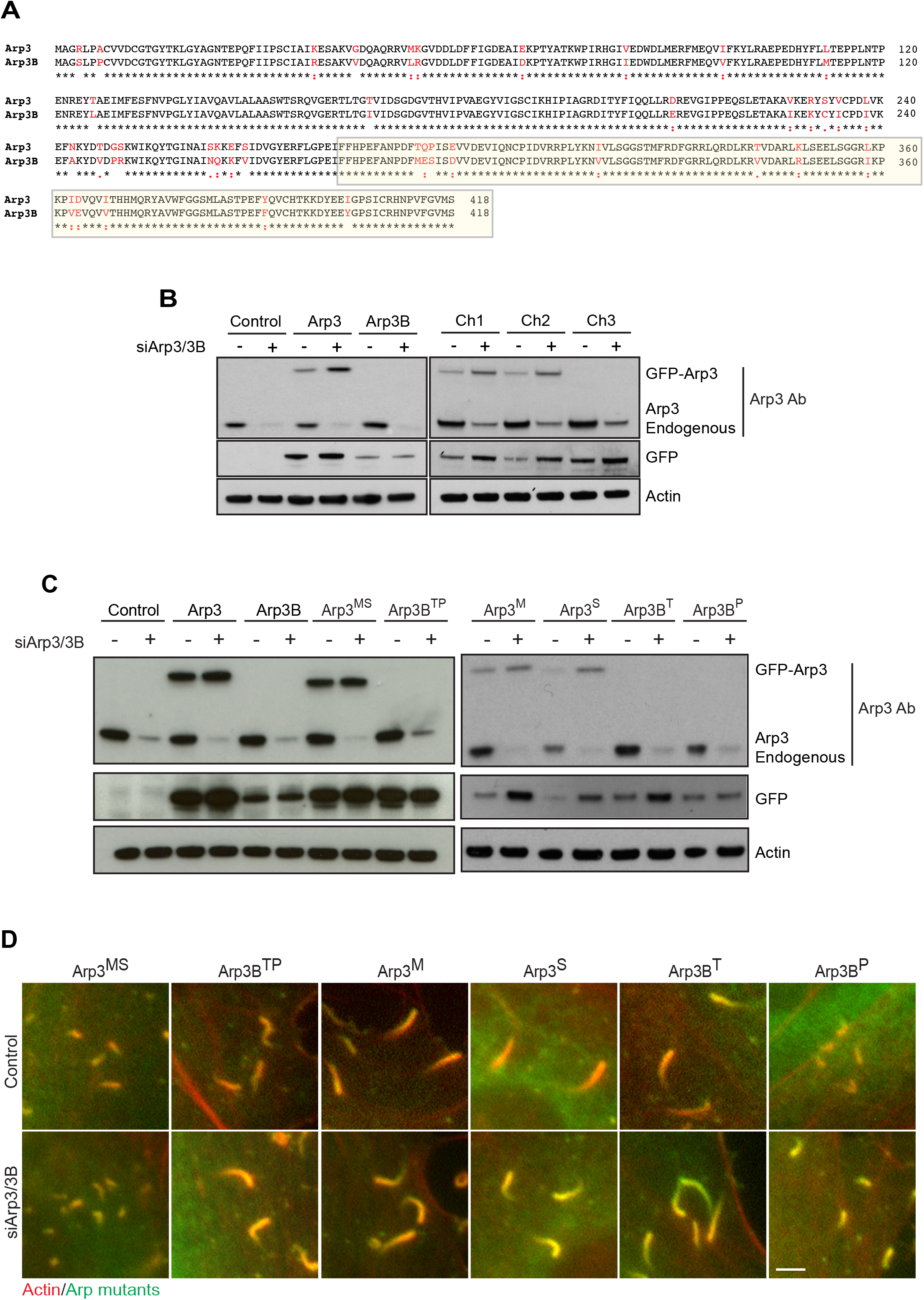
Arp3 and Arp3B chimeras and mutants. **A** Alignment of human Arp3 and Arp3B showing conservative residues (black) and nonconservative amino acid differences (red). The C-terminal third of Arp3/3B responsible for long or short actin tail phenotypes is boxed in yellow. **B** The immunoblot shows the expression level of RNAi resistant GFP-tagged Arp3 and Arp3B together with the three different chimeras in HeLa cells treated with Arp3 and Arp3B siRNA. **C** Immunoblot analysis of the expression level of the indicated RNAi resistant GFP-tagged Arp3/3B mutants in HeLa cells treated with Arp3 and Arp3B siRNA. **D** Immunofluorescence images of representative actin tails (red) in HeLa cells stably expressing the indicated GFP-tagged Arp3/3B mutants (green) and treated with the indicated siRNA. Scale bars = 5 μm.

**Figure S4.**
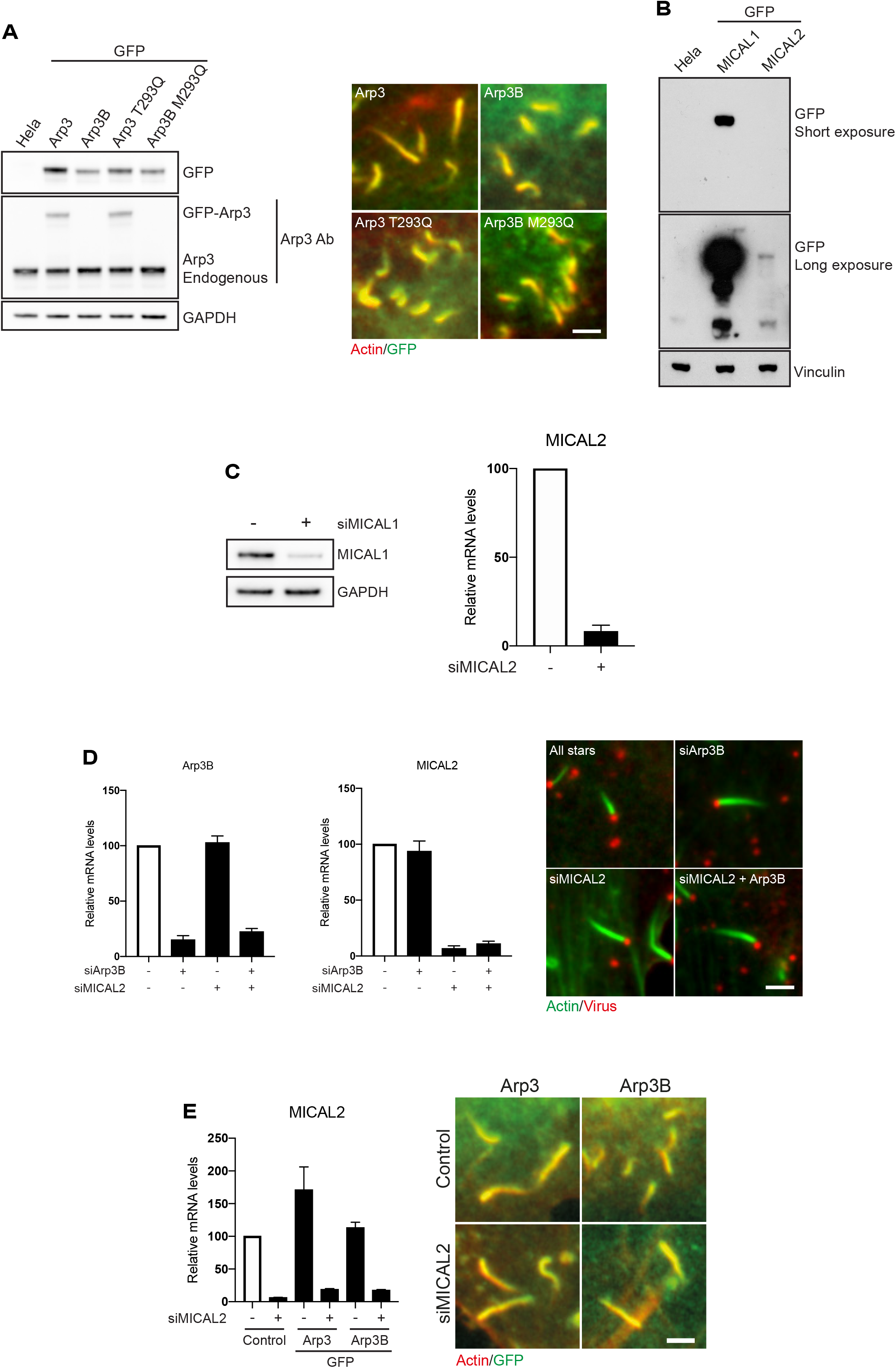
MICAL2 regulates actin tail length. **A** The immunoblot shows the expression level of GFP-tagged Arp3 and Arp3B together with their respective glutamine mutants. The immunofluorescence images show representative actin tails (red) in HeLa cells stably expressing the indicated GFP-tagged proteins (green). **B** Immunoblot analysis of the level of stable GFP-tagged MICAL1 and MICAL2 expression in HeLa cells. **C** Immunoblot and RT-PCR analysis of the level of MICAL1 protein or MICAL2 mRNA respectively in HeLa cells treated with MICAL1 and MICAL2 siRNA. **D** RT-PCR analysis of the level of Arp3B and MICAL2 mRNA in cells treated with Arp3B and MICAL2 siRNA. The images show representative actin tails (green) induced by vaccinia (red) in cells treated with the indicated siRNA. **E** RT-PCR analysis of the level of MICAL2 mRNA in cells treated with MICAL2 siRNA and stably expressing GFP-tagged Arp3 or Arp3B. The images show representative actin tails (red) in cells stably expressing GFP-tagged Arp3 or Arp3B (green) and treated with MICAL2 siRNA. Error bars represent SEM from three independent experiments. Scale bars = 2.5 μm.

**Figure S5.**
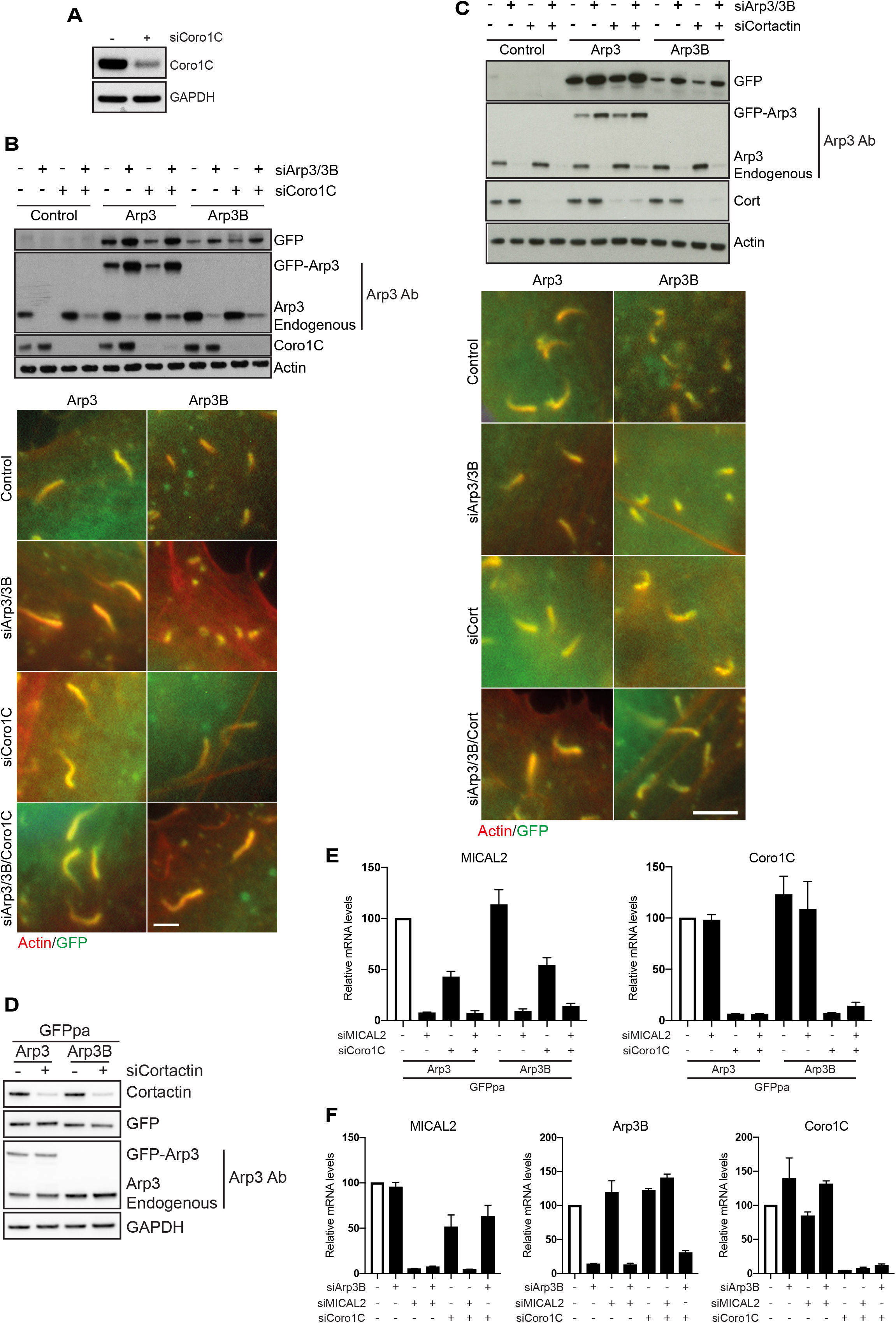
Coronin-1C and Cortactin recruit MICAL2 to actin networks. **A** Immunoblot analysis of the level of coronin-1C in HeLa cells treated with coronin-1C siRNA. **B** The immunoblot shows the expression level of RNAi resistant GFP-tagged Arp3 and Arp3B in HeLa cells treated with Arp3/Arp3B and coronin-1C siRNA. The immunofluorescent images show representative actin tails (red) in the treated HeLa cells stably expressing GFP-tagged Arp3 and Arp3B (green). **C** The immunoblot shows the expression level of RNAi resistant GFP-tagged Arp3 and Arp3B in HeLa cells treated Arp3/Arp3B and cortactin siRNA. The immunofluorescent images show representative actin tails (red) in the treated cells. Scale bar in B and C = 5 μm. **D** Immunoblot analysis of the level of cortactin in HeLa cells stably expressing GFP^PA^-tagged Arp3 and Arp3B and treated with cortactin siRNA. **E** RT-PCR analysis of the level of MICAL2 and coronin-1C mRNA in HeLa cells stably expressing GFP^PA^-tagged Arp3 and Arp3B and treated MICAL2 and/or coronin-1C siRNA. **F** RT-PCR analysis of the level of MICAL2, Arp3B and coronin-1C mRNA in HeLa cells stably expressing Cherry-GFP^PA^-β-actin and treated different combinations of MICAL2, Arp3B and coronin-1C siRNA.

### Movie legends

**Movie S1** The movie shows a time course of actin assembly (0.6 μM) in the presence of 2.5 nM Arp2/3 complex, containing ARPC1A and ARPC5 and activated by 50 nM GST-(Human WASp-VCA) acquired by total internal reflection fluorescence (TIRF) microscopy. The Arp3 isoform is indicated in each panel together with the time in seconds. The Arp3B data for the first 560 seconds is shown in Fig. 2B. Scale bar = 15 μm.

**Movie S2** The movie shows a representative example of photoactivation of Cherry-GFP^PA^-β-actin in a vaccinia actin tail in cells treated with control siRNA for 72 hours (also see Fig. S2C). HeLa cells stably expressing Cherry-GFP^PA^-β-actin (red before photoactivation and green after activation) were infected with Vaccinia expressing YFP-A3 for 8 hours. The time in seconds is indicated and the scale bar = 15 μm.

**Movie S3** The movie shows a representative example of photoactivation of Cherry-GFP^PA^-β-actin in a vaccinia actin tail in cells treated with Arp3B siRNA for 72 hours (also see Fig. S2E). HeLa cells stably expressing Cherry-GFP^PA^-β-actin (red before photoactivation and green after activation) were infected with Vaccinia expressing YFP-A3 for 8 hours. The time in seconds is indicated and the scale bar = 15 μm.

**Movie S4** The movie shows a representative example of photoactivation of GFP^PA^-Arp3 (appears green after activation) in a vaccinia actin tail, the analysis of which is presented in Fig. 2E and image stills in Fig. S2E. HeLa cells stably expressing GFP^PA^-Arp3 were infected with Vaccinia expressing YFP-A3 for 8 hours prior to imaging. The time in seconds is indicated and the scale bar = 15 μm.

**Movie S5** The movie shows a representative example of photoactivation of GFP^PA^-Arp3B (appears green after activation) in a vaccinia actin tail, the analysis of which is presented in Fig. 2E and image stills in Fig. S2E. HeLa cells stably expressing GFP^PA^-Arp3B were infected with Vaccinia expressing YFP-A3 for 8 hours prior to imaging. The time in seconds is indicated and the scale bar = 15 μm.

**Movie S6** The movie shows a representative example of the recruitment of GFP-MICAL2 (green) to a vaccinia induced actin tail visualized with LifeAct-iRFP670 (magenta) at 8 hours post infection. The time in seconds is indicated and the scale bar = 5 μm

**Movie S7** The movie shows a representative example of localization of GFP-MICAL1 (green) in a vaccinia infected HeLa cell at 8 hours post infection. Virus induced actin tails are visualized with LifeAct-iRFP670 (magenta). The time in seconds is indicated and the scale bar = 5 μm

**Movie S8** The movie shows a representative example of the recruitment of GFP-MICAL2 (green) to a vaccinia induced actin tails labelled with LifeAct-iRFP670 (magenta) at 8 hours post infection in cells treated with control siRNA for 72 hours. The time in seconds is indicated and the scale bar = 5 μm

**Movie S9** The movie showing the lack of GFP-MICAL2 (green) recruitment to vaccinia induced actin tails labelled with LifeAct-iRFP (magenta) in the absence of coronin-1C at 8 hours post infection. HeLa cells were treated with coronin-1C siRNA for 72 hours prior to infection. The time in seconds is indicated and the scale bar = 5 μm

## References

1. L. M. Machesky, S. J. Atkinson, C. Ampe, J. Vandekerckhove, T. D. Pollard, Purification of a cortical complex containing two unconventional actins from Acanthamoeba by affinity chromatography on profilin-agarose. J Cell Biol 127, 107–115 (1994).

2. E. D. Goley, M. D. Welch, The ARP2/3 complex: an actin nucleator comes of age. Nat Rev Mol Cell Biol 7, 713–726 (2006).

3. J. Muller et al., Sequence and comparative genomic analysis of actin-related proteins. Mol Biol Cell 16, 5736–5748 (2005).

4. J. L. Gordon, L. D. Sibley, Comparative genome analysis reveals a conserved family of actin-like proteins in apicomplexan parasites. BMC genomics 6, 179 (2005).

5. P. Jay et al., ARP3beta, the gene encoding a new human actin-related protein, is alternatively spliced and predominantly expressed in brain neuronal cells. Eur J Biochem 267, 2921–2928 (2000).

6. M. K. Balasubramanian, A. Feoktistova, D. McCollum, K. L. Gould, Fission yeast Sop2p: a novel and evolutionarily conserved protein that interacts with Arp3p and modulates profilin function. EMBO J 15, 6426–6437 (1996).

7. T. H. Millard, B. Behrendt, S. Launay, K. Futterer, L. M. Machesky, Identification and characterisation of a novel human isoform of Arp2/3 complex subunit p16-ARC/ARPC5. Cell Motil Cytoskeleton 54, 81–90 (2003).

8. W. H. Kahr et al., Loss of the Arp2/3 complex component ARPC1B causes platelet abnormalities and predisposes to inflammatory disease. Nature communications 8, 14816 (2017).

9. T. W. Kuijpers et al., Combined immunodeficiency with severe inflammation and allergy caused by ARPC1B deficiency. J Allergy Clin Immunol 140, 273–277 e210 (2017).

10. R. Somech et al., Disruption of Thrombocyte and T Lymphocyte Development by a Mutation in ARPC1B. J Immunol 199, 4036–4045 (2017).

11. I. Brigida et al., T-cell defects in patients with ARPC1B germline mutations account for combined immunodeficiency. Blood 132, 2362–2374 (2018).

12. S. Volpi et al., A combined immunodeficiency with severe infections, inflammation, and allergy caused by ARPC1B deficiency. J Allergy Clin Immunol 143, 2296–2299 (2019).

13. L. O. Randzavola et al., Loss of ARPC1B impairs cytotoxic T lymphocyte maintenance and cytolytic activity. J Clin Invest 129, 5600–5614 (2019).

14. W. Roman et al., Myofibril contraction and crosslinking drive nuclear movement to the periphery of skeletal muscle. Nat Cell Biol 19, 1189–1201 (2017).

15. J. V. Abella et al., Isoform diversity in the Arp2/3 complex determines actin filament dynamics. Nat Cell Biol 18, 76–86 (2016).

16. S. K. Donnelly, I. Weisswange, M. Zettl, M. Way, WIP Provides an Essential Link between Nck and N-WASP during Arp2/3-Dependent Actin Polymerization. Curr Biol 23, 999–1006 (2013).

17. F. Leite, M. Way, The role of signalling and the cytoskeleton during Vaccinia Virus egress. Virus Res 209, 87–99 (2015).

18. X. Snetkov, I. Weisswange, J. Pfanzelter, A. C. Humphries, M. Way, NPF motifs in the vaccinia virus protein A36 recruit intersectin-1 to promote Cdc42:N-WASP-mediated viral release from infected cells. Nat Microbiol 1, 16141 (2016).

19. A. Di Nardo et al., Arp2/3 complex-deficient mouse fibroblasts are viable and have normal leading-edge actin structure and function. Proc Natl Acad Sci U S A 102, 16263–16268 (2005).

20. A. Steffen et al., Filopodia formation in the absence of functional WAVE- and Arp2/3-complexes. Mol Biol Cell 17, 2581–2591 (2006).

21. D. J. Barry, G. A. Williams, C. Chan, Automated analysis of filamentous microbial morphology with AnaMorf. Biotechnol Prog 31, 849–852 (2015).

22. S. L. Liu, J. R. May, L. A. Helgeson, B. J. Nolen, Insertions within the actin core of actin-related protein 3 (Arp3) modulate branching nucleation by Arp2/3 complex. J Biol Chem 288, 487–497 (2013).

23. A. Zimmet et al., Cryo-EM structure of NPF-bound human Arp2/3 complex and activation mechanism. Sci Adv 6, eaaz7651 (2020).

24. M. Shaaban, S. Chowdhury, B. J. Nolen, Cryo-EM reveals the transition of Arp2/3 complex from inactive to nucleation-competent state. Nat Struct Mol Biol, (2020).

25. R. J. Hung, C. W. Pak, J. R. Terman, Direct redox regulation of F-actin assembly and disassembly by Mical. Science 334, 1710–1713 (2011).

26. S. Fremont, G. Romet-Lemonne, A. Houdusse, A. Echard, Emerging roles of MICAL family proteins - from actin oxidation to membrane trafficking during cytokinesis. J Cell Sci 130, 1509–1517 (2017).

27. D. J. Bigelow, T. C. Squier, Thioredoxin-dependent redox regulation of cellular signaling and stress response through reversible oxidation of methionines. Mol Biosyst 7, 2101–2109 (2011).

28. J. C. Aledo, Methionine in proteins: The Cinderella of the proteinogenic amino acids. Protein Sci 28, 1785–1796 (2019).

29. W. M. Brieher, H. Y. Kueh, B. A. Ballif, T. J. Mitchison, Rapid actin monomer-insensitive depolymerization of Listeria actin comet tails by cofilin, coronin, and Aip1. J Cell Biol 175, 315–324 (2006).

30. L. Cai, A. M. Makhov, D. A. Schafer, J. E. Bear, Coronin 1B antagonizes cortactin and remodels Arp2/3-containing actin branches in lamellipodia. Cell 134, 828–842 (2008).

31. N. A. Kulak, G. Pichler, I. Paron, N. Nagaraj, M. Mann, Minimal, encapsulated proteomic-sample processing applied to copy-number estimation in eukaryotic cells. Nat Methods 11, 319–324 (2014).

32. G. Hiller, K. Weber, Golgi-derived membranes that contain an acylated viral polypeptide are used for vaccinia virus envelopment. J. Virol. 55, 651–659 (1985).

33. S. Fremont et al., Oxidation of F-actin controls the terminal steps of cytokinesis. Nature communications 8, 14528 (2017).

34. S. S. Giridharan, J. L. Rohn, N. Naslavsky, S. Caplan, Differential regulation of actin microfilaments by human MICAL proteins. J Cell Sci 125, 614–624 (2012).

35. C. H. Durkin et al., RhoD Inhibits RhoC-ROCK-Dependent Cell Contraction via PAK6. Dev Cell 41, 315–329 e317 (2017).

36. R. C. Robinson et al., Crystal structure of Arp2/3 complex. Science 294, 1679–1684 (2001).

